# *Pseudomonas aeruginosa* ethanol oxidation by AdhA in low oxygen environments

**DOI:** 10.1101/670117

**Authors:** Alex W. Crocker, Colleen E. Harty, John H. Hammond, Sven D. Willger, Pedro Salazar, Nicholas J. Jacobs, Deborah A. Hogan

**Author notes:** To whom correspondence should be addressed Department of Microbiology and Immunology, Geisel School of Medicine at Dartmouth, Rm 208 Vail Building, Hanover, NH 03755.

## Abstract

*Pseudomonas aeruginosa* has a broad metabolic repertoire that facilitates its co-existence with different microbes. Many microbes secrete products that *P. aeruginosa* can then catabolize, including ethanol, a common fermentation product. Here, we show that under oxygen limiting conditions *P. aeruginosa* utilizes AdhA, an NAD-linked alcohol dehydrogenase, as a previously undescribed means for ethanol catabolism. In a rich medium containing ethanol, AdhA, but not the previously described PQQ-linked alcohol dehydrogenase, ExaA, oxidizes ethanol and leads to the accumulation of acetate in culture supernatants. AdhA-dependent acetate accumulation, and the accompanying decrease in pH, promotes *P. aeruginosa* survival in LB-grown stationary phase cultures. The transcription of *adhA* is elevated by hypoxia and in anoxic conditions, and we show that it is regulated by the Anr transcription factor. We have shown that *lasR* mutants have higher levels of Anr-regulated transcripts in low oxygen conditions compared to their wild type counterparts. Here, we show that a *lasR* mutant, when grown with ethanol, has an even larger decrease in pH than WT that is dependent on both *anr* and *adhA*. The large increase in AdhA activity similar to that of a strain expressing a hyperactive Anr-D149A variant. Ethanol catabolism in *P. aeruginosa* by AdhA supports growth on ethanol as a sole carbon source and electron donor in oxygen-limited settings and in cells growing by denitrification in anoxic conditions. This is the first demonstration of a physiological role for AdhA in ethanol oxidation in *P. aeruginosa*.

**Importance:** Ethanol is a common product of microbial fermentation, and the *Pseudomonas aeruginosa* response to and utilization of ethanol is relevant to our understanding of its role in microbial communities. Here, we report that the putative alcohol dehydrogenase, AdhA, is responsible for ethanol catabolism and acetate accumulation in low oxygen conditions and that it is regulated by Anr.

## Introduction

*Pseudomonas aeruginosa* has broad catabolic potential that contributes to its ability to thrive in diverse environments that often include other microbes. One environment in which *P. aeruginosa* metabolism has been well studied is in the mucus that accumulates during chronic infections in the lungs of individuals with cystic fibrosis. In this low-oxygen setting, *P. aeruginosa* encounters other pathogens, many of which are known to produce ethanol by fermentation (1). Ethanol has been detected in the volatiles expired from the lungs of CF patients infected with *P. aeruginosa* (2, 3), and several previous studies have shown that the addition of ethanol to the growth medium can alter *P. aeruginosa* metabolism, motility, and virulence factor production (4-7).

Ethanol metabolism in *P. aeruginosa* has mainly been studied with ethanol supplied as a sole carbon source under aerobic conditions, and that focus has been on the PQQ (pyrroloquinoline quinone)-dependent ethanol-oxidizing enzyme, ExaA, which donates electrons directly to the cytochrome 550, ExaB (8-11). Görisch and colleagues (8-10) described aerobic ethanol catabolism in detail, including a complex regulatory cascade initiated by ErbR/AgmR. Disruption of *exaA, exaB*, or genes involved in PQQ biosynthesis eliminates the ability to grow aerobically on ethanol as a sole carbon source on agar medium (11).

The *P. aeruginosa* genome encodes putative alcohol dehydrogenases other than ExaA but the roles of these others in the oxidation of ethanol has not been clearly established. The *P. aeruginosa* PAO1 gene encoded by PA5427 was named AdhA based on the high structural similarity of its crystal structure to diverse characterized alcohol dehydrogenase enzymes that use NAD+ as a cofactor (12). AdhA contains a catalytic domain (amino acids 1–157 and 292– 342) with a Zn+2 active site and a coenzyme-binding domain (residues 158–291) with a Rossmann fold motif that is characteristic of nicotinamide-adenine (NAD+) coenzyme-binding proteins (12, 13). The biological activity of this enzyme as an ethanol dehydrogenase in P. aeruginosa, however, was not described.

In *P. aeruginosa, adhA* transcripts are markedly elevated during hypoxic growth and during anoxic conditions in the presence of nitrate (14-17). Furthermore, *adhA* expression was greatly reduced upon deletion of *anr*, which encodes the oxygen-responsive transcription factor that controls the expression of genes involved in anaerobic and microoxic metabolism (14, 15). Schobert and colleagues (18) examined *P. aeruginosa* AdhA for the potential to participate in the conversion of pyruvate to produce ethanol when oxygen is limiting. Although lactic acid and acetic acid were formed, no evidence for ethanol formation was found.

In this work, we show that in rich medium amended with ethanol, ethanol was oxidized with the accumulation of acetate under oxygen limiting conditions and this was not dependent on ExaA but did require AdhA. We also found that acetate accumulation and the concomitant reduction in pH increased survival during stationary phase. The *adhA* mutant was also incapable of growth with ethanol as sole carbon source and electron donor in microoxic or anoxic conditions with nitrate as an electron acceptor. We provide evidence that the transcription of *adhA* is regulated by Anr in both LB medium and during growth on ethanol as a sole carbon source in low oxygen or anoxic environments. Together, these data suggest that AdhA is capable of ethanol oxidation for growth and for conversion to acetate and that it can work in conjunction with ExaAB to enable *P. aeruginosa* to catabolize ethanol in diverse environments.

## Results

### AdhA is responsible for accumulation of acetate from ethanol in LB

We have previously reported that *P. aeruginosa* strain PA14 grown in LB medium with 1% ethanol yields a lower final pH in stationary phase culture supernatants than cultures grown in LB alone (4). Due to the production of ammonia from amino acid catabolism, the culture pH from early stationary phase *P. aeruginosa* strain PA14 cultures (12 h, OD ∼4) was 1.5 pH units higher than uninoculated medium (pH 8.7 versus pH 7.2 for uninoculated medium). When ethanol was present in the medium, the final culture pH was lower than when cultures were grown in LB alone (pH 8.1 versus pH 8.7) (**Fig. 1A**). HPLC analysis of *P. aeruginosa* wild type (WT) culture supernatants at this time point (12 h) found that 36.3 mM ethanol had been consumed (**Fig. 1B**) and 11.1 mM acetate had accumulated in the medium (**Fig. 1C**). No acetate was detected in LB cultures without added ethanol and no lactate, pyruvate or succinate were detected in either the LB or LB + ethanol culture supernatants by the HPLC method used (data not shown). The final OD of cultures grown in LB + 1% ethanol were similar for cultures grown with and without ethanol. All strains yielded a slightly higher OD_600_ value in the presence of ethanol, but the difference did not reach statistical significance for the wild type (**Fig. 1D**).

**Figure 1.**
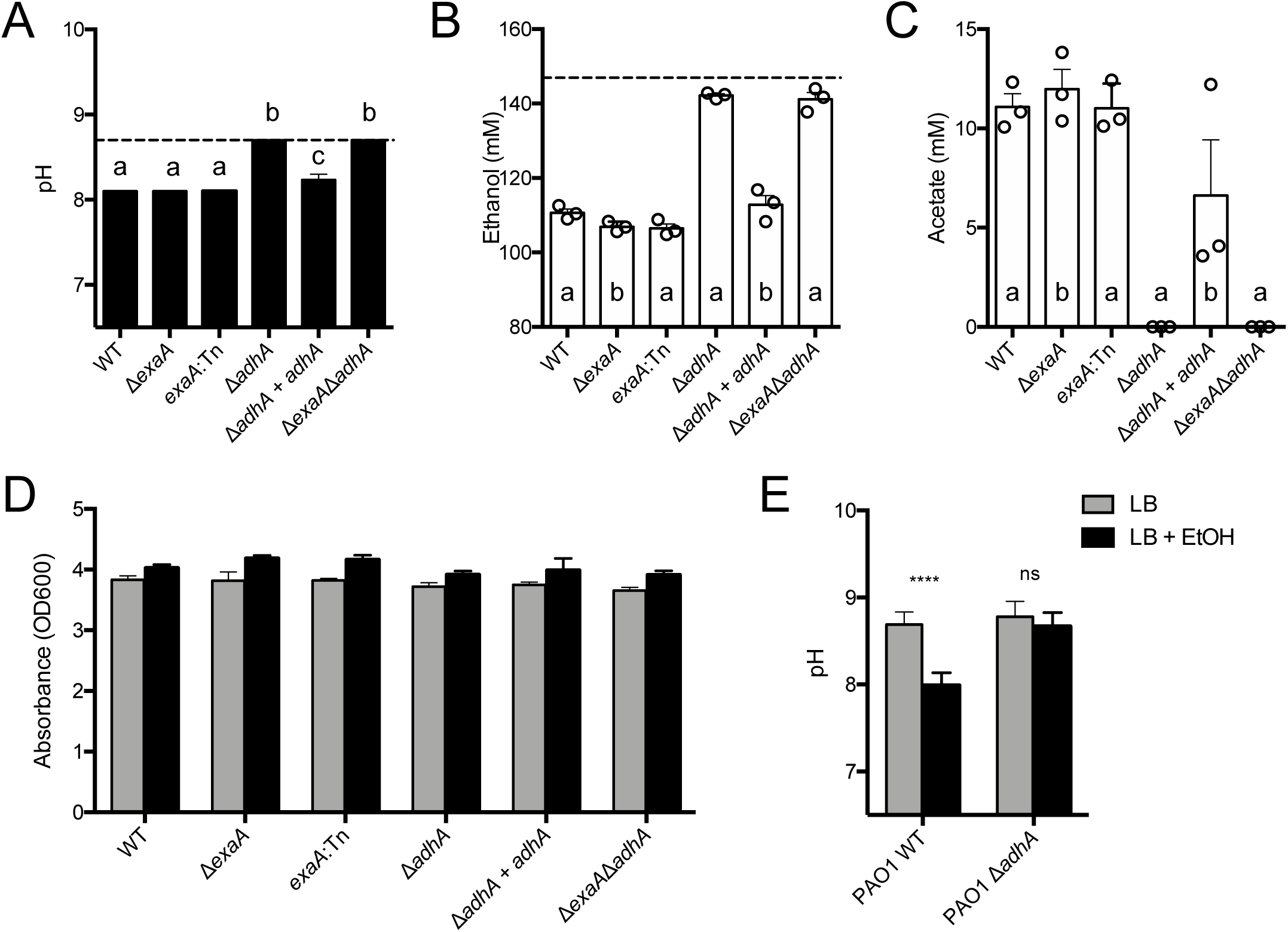
Ethanol catabolism by *P. aeruginosa* leads to lower medium pH through the accumulation acetate. *P. aeruginosa* PA14 strains including wild type, Δ*exaA, exaA*::Tn*M*, Δ*adhA*, Δ*adhA*+*adhA*, nd Δ*exaA*Δ*adhA* were grown in LB for 12 h in tubes on a roller drum. A. pH of cultures, measured sing pH indicator strips, in LB (dotted line) and LB+ ethanol (bars). B. Ethanol remaining in culture pernatants relative to uninoculated LB + 1% ethanol (dotted line). C. Acetate concentrations in pernatants in LB + 1% ethanol in the same cultures analyzed in panels A and B. D. Optical ensity of strains grown in LB (grey) and LB+1% ethanol (black). E. *P. aeruginosa* strain PAO1 wild pe and Δ*adhA* final culture pH values after 16 h of growth in LB (grey) or LB with 1% ethanol lack). All cultures were grown at 37°C. In panels A-C, samples with different letters are significantly fferent, *P*<0.05. In D, ethanol-grown cultures were significantly higher (*P*<0.05) for each strain xcept WT and Δ*adhA*. In E, **** indicates a *P*<0.001, ns is not significant.

To determine if ethanol catabolism in LB was due to activity of the characterized ethanol dehydrogenase ExaAB, we examined ethanol catabolism in a mutant with an in-frame deletion of *exaA* and a validated *exaA*::Tn*M* insertion mutant (5, 19), both of which were generated in the strain PA14 background. Both mutants still acidified culture medium, consumed ethanol, and accumulated acetate in cultures with ethanol present, suggesting that ExaA was not required and that another ethanol catabolic enzyme was present (**Fig. 1A-C**). In light of this result, we examined whether AdhA, a putative NAD+-linked alcohol dehydrogenase encoded by PA14_71630 (PA5427 in strain PAO1) contributed to ethanol catabolism and acetate accumulation. We constructed and analyzed an in-frame mutant lacking *adhA* and found that it did not have a lower pH when grown in LB + 1% ethanol compared to LB alone (**Fig. 1A**). The Δ*adhA* mutant showed only a minor decrease in the ethanol concentration of the medium (4.7 mM), and no acetate was detected (**Fig. 1B and C**). The defect in the *adhA* mutant could be complemented by replacement of the *adhA* gene at the native locus (**Figs. 1A-C**), while a Δ*adhA*Δ*exaA* double mutant phenocopied the Δ*adhA* mutant. Together, these data indicate that ethanol was required for accumulation of acetate in *P. aeruginosa* cultures grown in LB, and that this phenomenon was dependent on *adhA*, but not *exaA*, in *P. aeruginosa* strain PA14. A similar requirement for *adhA* for acidification of LB medium in cultures with ethanol was observed in *P. aeruginosa* strain PAO1, a genetically distinct laboratory strain (**Fig. 1E**).

### Ethanol catabolism through AdhA increases stationary phase survival

Based on our initial observations that cultures grown with ethanol remained turbid for longer in stationary phase, and on previous studies in *E. coli* that show ethanol can promote stationary phase survival in rich media such as LB (20), we tested the hypothesis that ethanol catabolism by AdhA promoted stationary phase survival. In LB cultures with no ethanol, strain PA14 wild type, Δ*adhA*, and Δ*adhA*Δ*exaA* all showed a significant decrease in viability over time (8h, 24h, and 48h) with no significant differences between strains (**Fig. 2-LB**). In LB with 1% ethanol, the wild-type cultures did not show a decrease in CFUs over time, indicating that ethanol was protective, but the Δ*adhA* and Δ*adhA*Δ*exaA* mutants did not receive a benefit from the presence of ethanol in the medium (**Fig. 2-LB+EtOH**). Consistent with our observation that AdhA activity was responsible for the lower culture pH in LB with ethanol at 12 h (**Fig. 1**), culture pH in LB with ethanol at 24 h showed even greater differences between wild type and the *adhA* or *adhA exaA* mutants (pH 6.5 for wild type and 8.7 for mutants). To determine if ethanol was protective in late stationary phase because of its effects on culture pH or due to other factors, the experiment also included cultures grown in LB that was buffered to pH 7. We found that maintaining a pH of 7 was sufficient to mitigate the loss of viability in late stationary phase relative to unbuffered medium in the wild type and all mutant strains (**Fig. 2-LB pH 7 versus LB**) similar to what has been observed for *E. coli* (21). Together, these data suggest that the ability of AdhA to convert ethanol to acetate in the late stationary phase of growth even in rich media can mitigate toxic alkaline pH conditions.

**Fig. 2.**
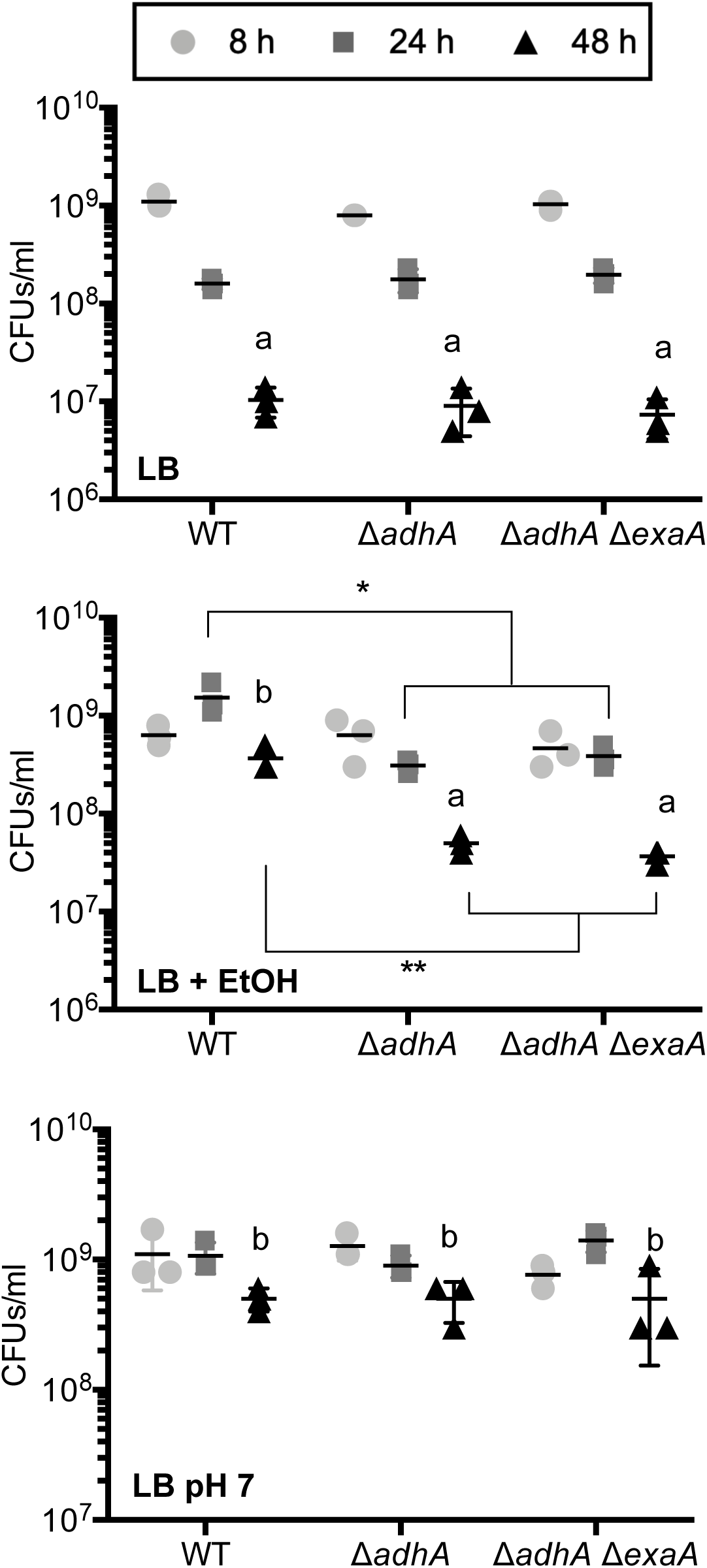
Ethanol and pH effects on *P. aeruginosa* stationary phase survival in LB. The CFUs from 8 h, h, and 48 h cultures were measured for PA14 wild type (WT), Δ*adhA*, and Δ*exaA*Δ*adhA* in LB p), LB with 1% ethanol (EtOH) (middle), and LB buffered to pH 7 (bottom) in a single experiment own in three panels for ease of comparison. * indicates *P*<0.05 and ** indicates *P*<0.01. Letters dicate significant differences between 48 h cultures across the three conditions tested with a-b *P*< 05. Data shown are from a single experiment with three independent replicates at each time point. e experiment was repeated with similar results two additional times.

### Anr regulates AdhA-dependent ethanol oxidation in wild type and Δ*lasR* strains

We and others have shown previously that *adhA* transcript levels are as much as 40-fold lower in a *Δanr* mutant, when compared to the wild type, during growth in microoxic or anoxic conditions (14, 15). Thus, we speculated that Anr was regulating *adhA* in dense LB cultures when oxygen levels decreased due to cell respiration. Indeed, like the Δ*adhA* strain, the *Δanr* mutant culture had a higher pH after growth in LB + 1% ethanol after 16 h than did the wild type or the *adhA* complemented strain, though the defect was not quite as striking as in the *adhA* strain (**Fig. 3**). We have previously published the creation of a strain in the PAO1 background in which native *anr* is replaced with an allele that encodes Anr-D149A, a variant that is active even in the presence of oxygen and demonstrated that the activity is 2-4 times higher than native Anr in microoxic environments (22). We constructed a PA14 strain in which *anr* was replaced with *anr-D149A*, and saw greater acidification of LB + 1% ethanol medium than the otherwise isogenic wild-type PA14 indicating that increased Anr activity leads to even more ethanol catabolism and acetate accumulation in LB medium (**Fig. 3**).

**Figure 3.**
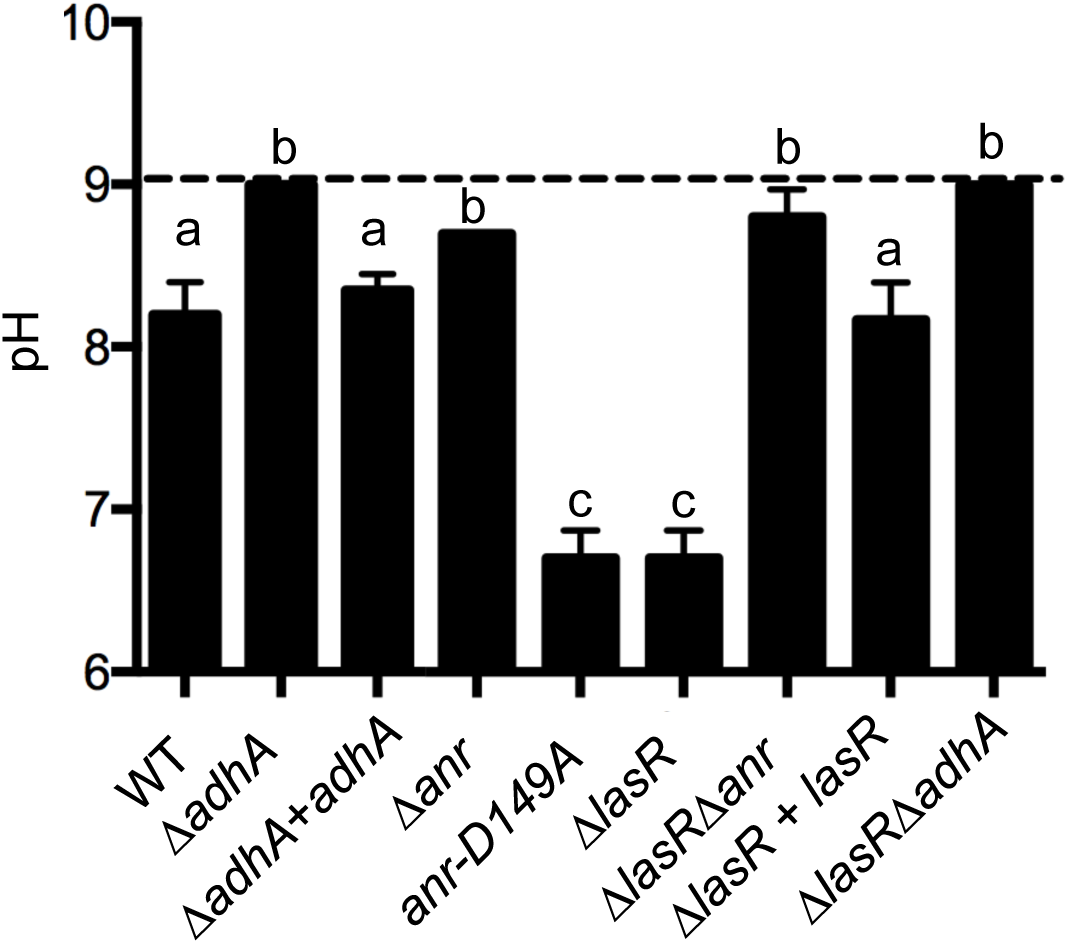
Culture pH after growth in LB + 1% ethanol for 16 h. The pH values for cultures of *P. aeruginosa* strain PA14 wild type (WT), Δ*adhA*, Δ*adhA*+*adhA*, Δ*anr, anr-D149A, ΔlasR, ΔlasRΔanr, ΔlasRΔadhA, ΔlasR+lasR* after growth in LB (dashed line) or LB with 1% ethanol (bars). Significance for comparisons of the LB+ethanol cultures was performed using a one-way ANOVA and Tukey’s multiple comparisons test; samples with the same letter were not significantly different (*P*>0.05).

*P. aeruginosa* isolates with loss-of-function mutations in *lasR* are commonly recovered from both the clinic and in the environment (23-26), and previous studies have shown that under microoxic conditions Anr activity is higher in both engineered and naturally-occurring *lasR*-defective strains relative to their counterparts with functional LasR (14, 23). In Hammond et al. (14), *adhA* was found to be five-fold higher (*P* <0.005) in *lasR* mutants compared to their genetically similar counterparts with *lasR* intact across four pairs of strains including strain PA14 and PA14 Δ*lasR*. Thus, we tested whether ethanol-dependent acidification of the medium was greater in Δ*lasR* mutant cultures. Like the strain bearing the Anr-D149A variant, Δ*lasR* cultures showed greater acidification at 16 hours compared to the wild type or the *lasR* complemented strain (**Fig. 3**). Deletion of either *adhA* or *anr* in the Δ*lasR* background eliminated the acidification phenotype (**Fig. 3**). Together, these data strongly suggest that Anr induction of AdhA promotes ethanol oxidation to acetate, resulting in reduced medium pH and that conditions or backgrounds that increase Anr activity will enhance the rate of ethanol conversion to acetate in LB.

### Anr-regulated AdhA can support growth of *P. aeruginosa* strains PA14 and PAO1 on ethanol as a sole carbon source in low oxygen and anoxic conditions

In light of *adhA* regulation by Anr (see **Fig. 3**) and its ability to oxidize ethanol to acetate in LB (see **Fig. 1B**), we sought to determine if AdhA would be required to support growth on ethanol as a sole carbon source in low oxygen environments. The Δ*adhA* mutant was grown in microoxic conditions (a 0.2% oxygen atmosphere) in M63 liquid medium with 1% ethanol as the sole carbon source. Under these low oxygen conditions, the Δ*adhA* strain showed a marked defect in growth that could be fully complemented by reconstitution of the mutant with the native *adhA* gene (**Fig. 4A**). The Δ*exaA* mutant did not have a defect in microoxic growth, and in fact showed a slight, significant increase in growth that could be complemented by the reconstitution of the *exaA* gene at the native locus. The Δ*exaA*Δ*adhA* mutant was similar in growth to the *adhA* single mutant (**Fig. 4A**). All strains grew equally well in M63 medium with glucose as the sole carbon source in 0.2% oxygen with an average OD_600_ of 0.71 +/- 0.09 across strains.

**Figure 4.**
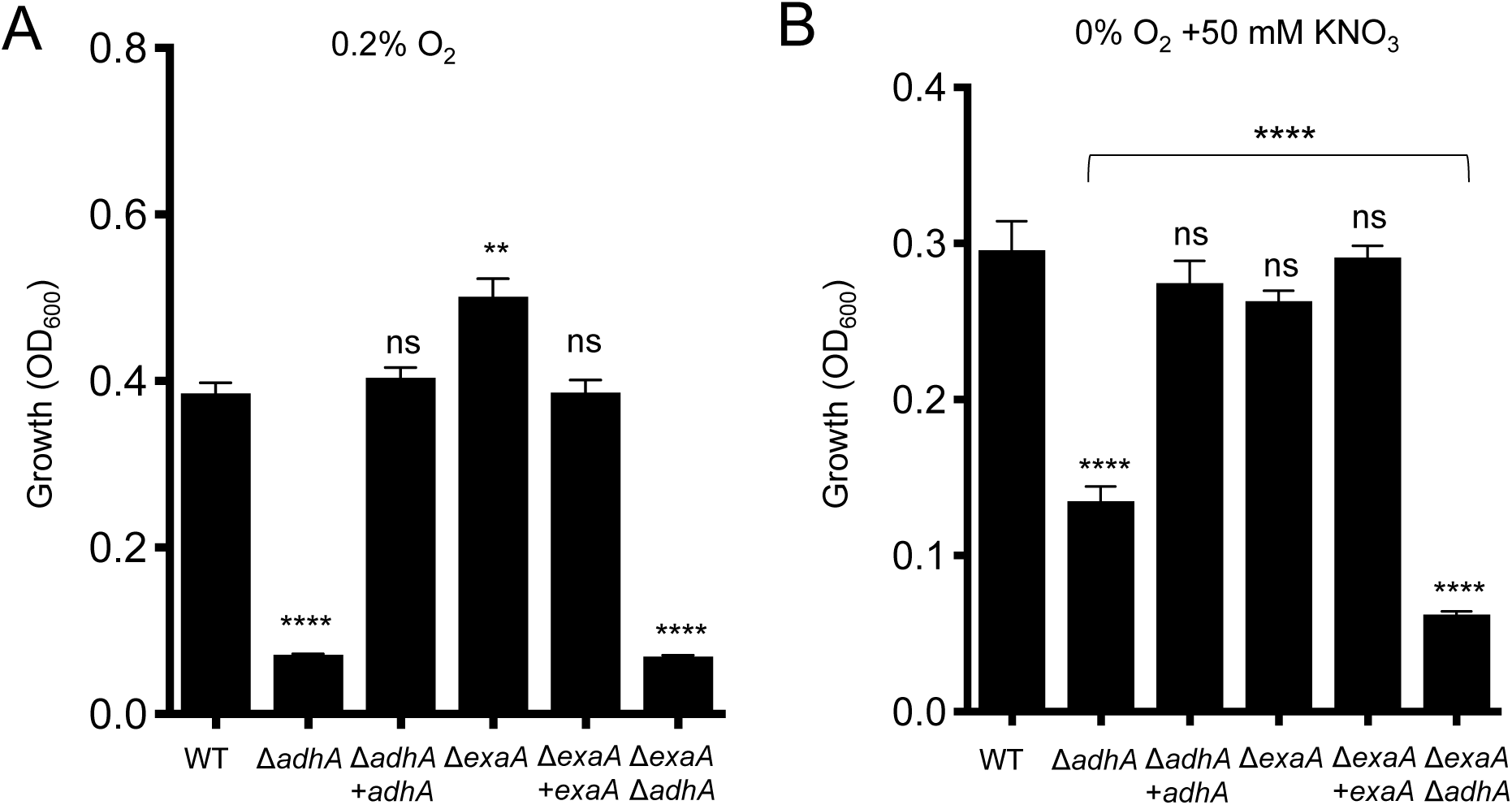
Growth of *P. aeruginosa* PA14 strains on ethanol as a sole carbon source. A. Growth in M63 liquid medium with ethanol as the sole carbon source in 0.2% oxygen. B. *P. aeruginosa* strains grown in M63 liquid medium with ethanol as a sole carbon source in anoxia with 50 mM KNO_3_. Data are representative of at least three independent experiments, each with three or more biological replicates. Statistics based on one-way ANOVA and Tukey’s multiple comparison; **, *P*<0.01, ****, *P*<0.001, ns, not significant; statistics presented indicate difference from the WT unless otherwise indicated.

AdhA could also support ethanol utilization as sole carbon source in *P. aeruginosa* when growing anoxically using nitrate respiration for energy generation (**Fig. 4B**). Wild type, the Δ*adhA* mutant, and the Δ*adhA* complemented strain were grown on M63 liquid medium with 1% ethanol as the sole carbon source and 50 mM nitrate. This experiment showed that the *adhA* mutant was essential for growth on ethanol in anoxic conditions. Although the delta exaA deletion grew quite well, the slightly reduced growth of the Δ*exaA* strain relative to the wild type and the Δ*exaA*Δ*adhA* double mutant relative to the Δ*adhA* single mutant did suggest that *exaA* might contribute slightly to ethanol utilization as sole carbon source under anoxic denitrifying conditions (**Fig. 4B**). Further study would be needed to evaluate this possibility. In anoxic conditions in medium with glucose as the sole carbon source and with nitrate as the terminal electron acceptor, all strains grew similarly to wild type with an OD_600_ of 0.39 +/- 0.05, except for the Δ*anr* mutant which did not grow. Our finding that AdhA is needed for growth on ethanol as sole carbon source under anoxic denitrifying conditions is consistent with observations that ethanol can be utilized as an electron donor under this same condition. Gliese *et al.* discussed unpublished results that an NAD(P) linked ethanol dehydrogenase was expressed under these denitrifying conditions (29), but no data or details about the exact nature of this ethanol dehydrogenase were reported in that paper.

Like Δ*adhA*, the *Δanr* mutant was unable to grow on ethanol as a sole carbon source at 0.2% oxygen, and that defect could be complemented by restoring *anr* to the strain (**Fig. 5**). To determine if the defect in growth in the Δ*anr* mutant on ethanol in low oxygen was due to the inability to induce *adhA*, a Δ*anr* strain expressing *adhA* under the control of an arabinose-inducible promoter was constructed. In this background, overexpression of *adhA* was sufficient to partially rescue growth of a Δ*anr* mutant relative to a Δ*anr* strain bearing the empty vector (**Fig. 5**). The inability to completely restore growth to the Δ*anr* mutant upon overexpression of *adhA* suggests that other Anr-regulated factors such as the high affinity cytochrome C oxidases are required for fitness in low oxygen environments.

**Figure 5.**
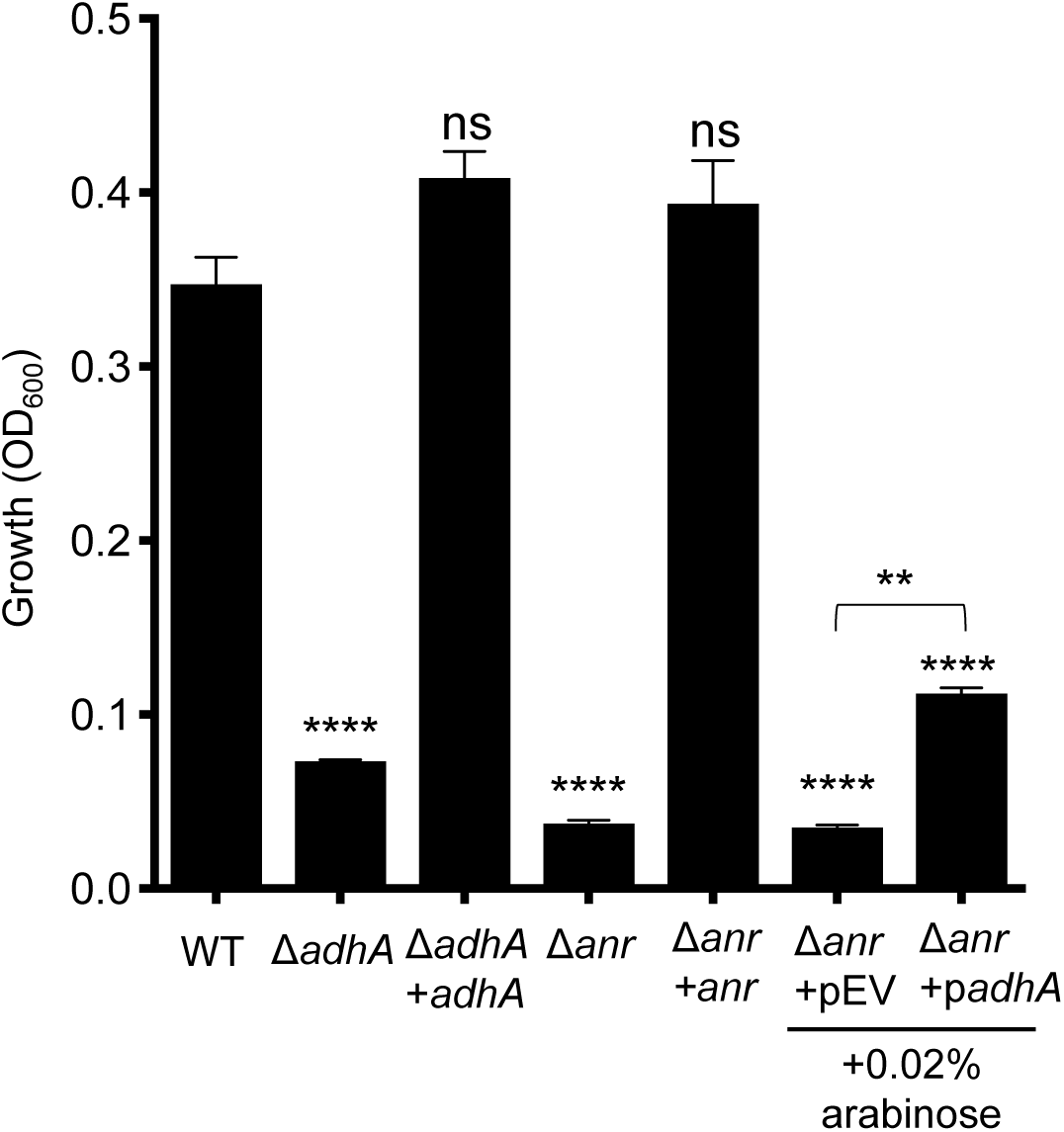
Anr regulation of *adhA* expression. *P. aeruginosa* strains grown in M63 liquid medium with ethanol as the sole carbon source in a 0.2% oxygen atmosphere. The strains included wild type strain PA14 (WT), Δ*adhA*, Δ*adhA*+*adhA*, Δ*anr*, Δ*anr*+*anr*, Δ*anr* containing an empty vector (pEV) or a plasmid with *adhA* under the control of an arabinose inducible promoter (p*adhA*). Statistics based on one-way ANOVA and Tukey’s multiple comparisons; unless otherwise indicated, statistical comparisons are to the WT culture; \*\*\*\**P*<0.0001; \*\**P*<0.01.

To complement our analysis of the role of AdhA in ethanol catabolism in liquid M63 with ethanol as the sole carbon source in microoxic and anoxic conditions, we also analyzed the role of AdhA in growth on ethanol in normoxic cultures conditions. Consistent with previously published results (5, 9, 10) *exaA* was necessary and sufficient for growth on ethanol on agar medium at 21% oxygen in both strains PA14 and PAO1. The *adhA* mutants in both PA14 and PAO1 showed no defect in aerobic growth on ethanol on an agar medium and the phenotype of the Δ*adhA*Δ*exaA* mutant was similar to that of the Δ*exaA* single mutant (**Fig. S1A**). However, in liquid medium (M63 with ethanol as the sole carbon source in 5 ml cultures in test tubes on a roller drum) both *exaA* and *adhA* were required for growth equal to that in the PA14 and PAO1 backgrounds (**Fig. S1 B and C, respectively**). These data suggest that even in normoxic atmospheric conditions, metabolic activity in cells within culture tubes created sufficient oxygen restriction to induce *adhA* under these conditions. All strains grew equally well in M63 medium with glucose as the sole carbon source with an average OD_600_ of 0.94 +/- 0.2. Together these data suggest that AdhA, under the control of Anr, can contribute to ethanol catabolism in diverse culture conditions, and in some cases can work in parallel with the ExaAB ethanol catabolic complex.

## Discussion

*Pseudomonas aeruginosa* is remarkable in its ability to grow in diverse environments. Although for years only one pathway for ethanol oxidation in this organism has been known (10), it is unsurprising that *P. aeruginosa* possesses other enzymes that carry out the same metabolic process, regulated by entirely different signals. Through this work, we show that AdhA contributes to the catabolism of ethanol in a rich medium (LB + 1% ethanol) and in a minimal medium with ethanol as the sole carbon source under oxygen-limiting conditions. In the absence of oxygen, ethanol supported growth as an electron donor for nitrate, a phenomenon also observed in the complex microbial communities of wastewater treatment systems (27). AdhA-dependent ethanol catabolism in LB was concomitant with acetate accumulation in culture supernatants, and the acidification that resulted increased *P. aeruginosa* survival in LB protecting against the excessive rise in pH due to deamination of amino acids. In light of studies in *E. coli*, acetate may be protective in stationary phase through other mechanisms as well (28). Together, these data suggest that ethanol is at times an important resource in oxygen-limited conditions.

Our data, in conjunction with other published transcriptomics analyses of the Anr regulon and gene expression comparisons between oxic and microoxic or anoxic environments (14-16, 29), showed that Anr, the oxygen responsive transcription factor, is a key regulator of *adhA.* It is perhaps curious that an NAD+-linked ethanol oxidizing system is activated by Anr and hypoxia since low oxygen conditions limit the oxidation of NADH. We, like others (18), have considered the possibility that *P. aeruginosa* uses AdhA to ferment pyruvate to ethanol, regenerating NAD+, but we never observed ethanol in supernatants from LB-grown cultures even when extracellular acetate was present. The regulation of *adhA* by Anr may be indicative of *P. aeruginosa* utilization of fermentation products, such as ethanol, as significant carbon sources in low oxygen or anoxic settings.

The finding that *P. aeruginosa* accumulates significant amounts of acetic acid from added ethanol has not been reported previously. Acetate accumulation in other species such as *Acetobacter sp.* and *Gluconobacter* spp., which are used for the commercial production of vinegar, generate large amounts of extracellular acetic acid from ethanol oxidation using an ExaAB like system including a PQQ-linked alcohol dehydrogenase (30, 31). A very early study reported that many *P. fluorescens* strains accumulated acetic acid from ethanol in a peptone medium, but in that study the *P. aeruginosa* strain did not, although it did use ethanol as a sole carbon source (32).

The question arises as to why this AdhA-dependent accumulation of acetate from ethanol had not been previously observed in *P. aeruginosa*. Previous studies largely investigated the aerobic oxidation and utilization of ethanol as a sole carbon source. In the present study, we observed acetate accumulation when we added ethanol to LB, a rich medium, and incubated it with *Pseudomonas* in a test tube on a roller drum apparatus. We hypothesize that the rapid respiratory activity of cells after reaching late log phase had depleted dissolved oxygen in the broth on the roller tube apparatus. This would be expected to increase the level of AdhA activity, which is markedly elevated by hypoxia as explained above.

The obvious question that arises is why the acetate formed was not further metabolized in the LB medium conditions. We hypothesize that this is due to the presence of amino acids in the medium activating the previously characterized catabolite repression of acetate utilization in *P. aeruginosa* (9, 33). The effects of both ambient oxygen and catabolite-repressing substrates on ethanol oxidation suggest that effects of ethanol on *P. aeruginosa* in communities with ethanol-producing taxa is modulated by a variety of environmental conditions.

## Materials and Methods

### Strains and growth conditions

Bacterial strains and plasmids used in this study are listed in **Table S1**. Bacteria were maintained on 1.5% agar LB (lysogeny broth) plates (34). Where stated, ethanol (200-proof) was added to the medium (liquid or molten agar) to a final concentration of 1%. For LB medium buffered to pH 7, 100mM HEPES acid was added to LB and the pH brought up to 7 by titration with NaOH. The *exaA*::Tn*M* was maintained on LB with 60 µg/mL gentamicin (19).

### Construction of in-frame deletions, complementation, and plasmids

Construction of plasmids, including in-frame deletion and complementation constructs, was completed using yeast cloning techniques in *Saccharomyces cerevisiae* as previously described (35) unless otherwise stated. Primers used for plasmid construction are listed in Table S2. In-frame deletion and single copy complementation constructs were made using the allelic replacement vector pMQ30 (35). Promoter fusion constructs were made using a modified pMQ30 vector with *lacZ*-*GFP* fusion integrating at the neutral *att* site on the chromosome. All plasmids were purified from yeast using Zymoprep™ Yeast Plasmid Miniprep II according to manufacturer’s protocol and transformed into electrocompetent *E. coli* strain S17 by electroporation. Plasmids were introduced into *P. aeruginosa* by conjugation and recombinants were obtained using sucrose counter-selection and genotype screening by PCR. The primers are listed in Table S2. The *exaA*::Tn*M* strain was confirmed with locus specific primers as previously published (5).

### Growth on ethanol as a sole carbon source

For the assessment of growth on ethanol as a sole carbon source, M63 medium with 1% ethanol was used (36). Microoxic cultures were grown in 1 mL medium in a single well of a 12-well plastic culture plate in a chamber with a controlled atmosphere (Coy Products) at 37°C set to maintain a 0.2% O_2_ atmosphere. M63 minimal medium with either 0.2% glucose or 1% ethanol as sole carbon sources was used. Cultures were inoculated at an initial OD_600_ of 0.05 into fresh medium from overnight liquid cultures grown in LB. OD_600_ was measured at 48 h post-inoculation using a Genesys 6 spectrophotometer. Normoxic cultures were grown in 5 mL of M63 minimal medium with either 0.2% glucose or 1% ethanol as sole carbon sources inoculated to an initial OD_600_ of 0.05 into fresh medium from overnight liquid cultures grown in LB in 18 × 150 mm borosilicate culture tubes at 37°C on a roller drum. OD_600_ was measured at 24 h post-inoculation using a Genesys 6 spectrophotometer. Anoxic cultures were grown in 2 mL of M63 minimal medium with either 0.2% glucose or 1% ethanol as sole carbon sources and supplemented with 50 mM KNO_3_. Cultures were inoculated to an initial OD_600_ of 0.05 into fresh medium from overnight liquid cultures grown in LB in 18mm borosilicate culture tubes at 37°C on a roller drum in 15 mL conical screw-cap tubes (Sarstedt). Tubes were placed in 2.5 liter AnaeroPack System rectangular jars (Mitsubishi Gas Chemical Co., Inc.) with one GasPak EZ sachet (BD). These cultures were grown for 5 days at 37°C prior to measuring OD_600_ using a Genesys 6 spectrophotometer.

### Ethanol catabolism experiments in LB medium

For experiments in which *P. aeruginosa* was grown in liquid LB, LB + 1% ethanol, or LB buffered to pH 7, strains were first grown overnight in LB for 14 hours, and 100 μL of the overnight culture was used to inoculate 5 mL of the test medium. For pH measurements, culture supernatant was applied to Millipore pH-indicator strips (pH 6.5 - 10.0); colors were compared to reference and values were recorded immediately.

### HPLC analysis of culture supernatants

For HPLC, culture supernatants were collected after pelleting cells by centrifugation for one minute at 13,000rpm. 400 μL of culture supernatant and 40 μL of 10% sulphuric acid solution were added to Costar Spin-X 0.22um filter centrifuge tubes and spun for 5 minutes at 13,000 rpm. 100 µL of each sample was transferred to an HPLC auto-sampler vial (Chemglass CV-1042-1232) and one 40 μL injection of each sample was run for 30 min. Chemical peaks were detected by refractive index using an Aminex HPX-87H column (Bio-Rad, Hercules, CA) with a 2.5 mM sulfuric acid solution mobile phase. Chemical concentrations were determined using calibrations curves based on three dilution standards of pyruvate, lactate, acetate, and ethanol.

### Statistics

Unless otherwise stated, data are based on three biological replicates with the mean and standard deviations calculated, and are representative of at least three independent experiments containing multiple replicates. Unless stated otherwise, means and standard deviations were calculated in Graph Pad Prism 8 and analyses were completed using a one-way ANOVA and Tukey’s multiple comparisons test, with *P*-values indicated in figure legends.

## Acknowledgements

Research reported in this publication was supported by National Institutes of Health (NIH) grant R01 AI 091702 and to D.A.H., NIAID T32 AI007519 to C.E.H., DNA Sequencing is supported in part by a Cancer Center Core Grant (P30 CA023108) from the National Cancer Institute. Support for the project was also provided by the NIGMS P20GM113132 through the Molecular Interactions and Imaging Core (MIIC) and the CF RDP STANTO15R0 and P30-DK117469 for statistical support and CFF (D.A.H.). The content of this publication is solely the responsibility of the authors and does not necessarily represent the official views of the NIH. We thank Judy Jacobs for important insights, literature reviews, and comments on the manuscript. We thank Daniel G. Olson for invaluable assistance with HPLC.

---

**Figure S1.**
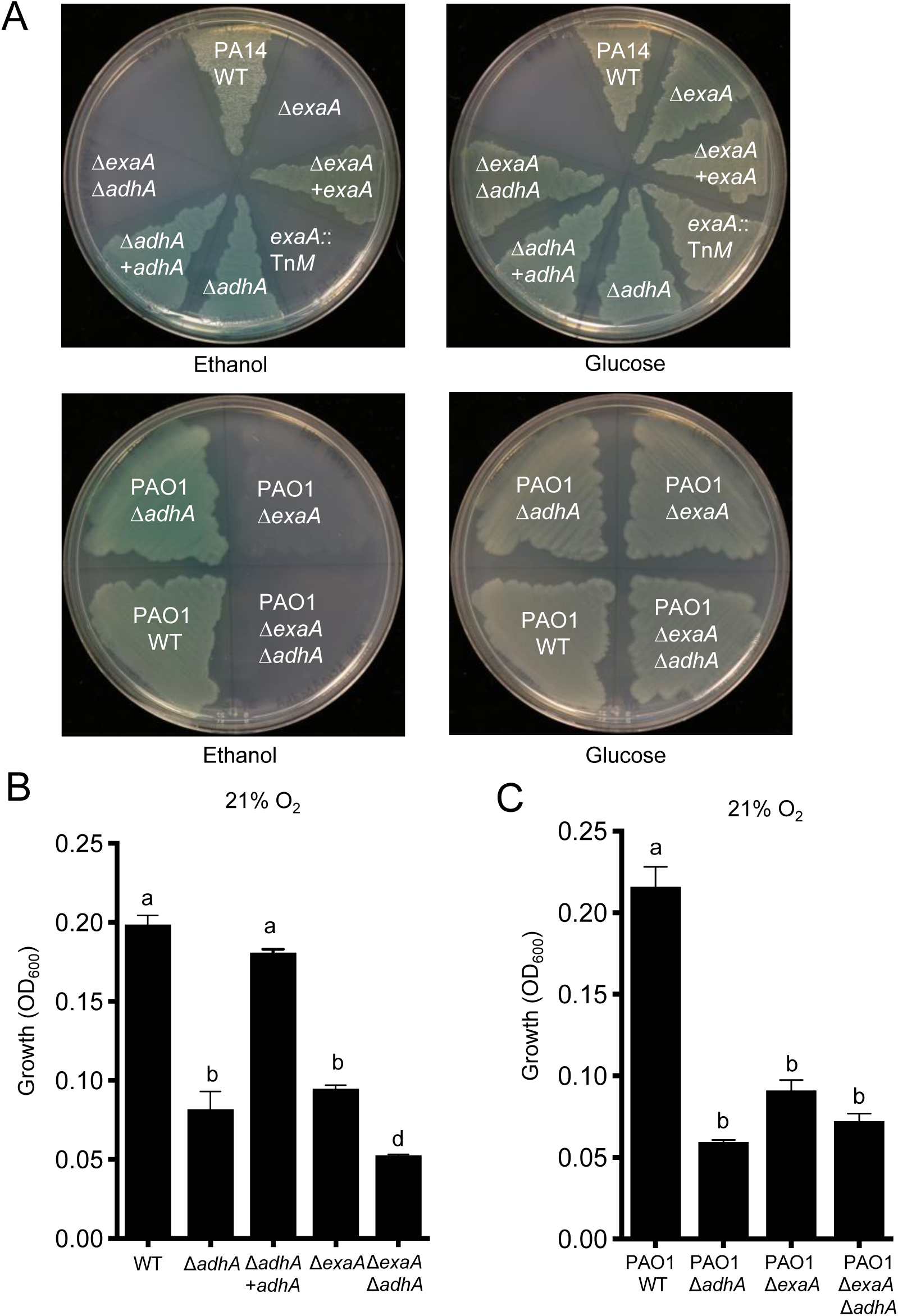
Growth of *P. aeruginosa* strains on ethanol as a sole carbon source. **A**. *P. aeruginosa* PA14 (top) and PAO1 (bottom) wild types and their *exaA* or *adhA* derivative and mutants after complementation. In each strain background, an Δ*exaA*Δ*adhA* strain is also shown. All strains were grown on 1.5% agar M63 with either ethanol or glucose, as indicated, as the sole carbon source at 37 C for 24 h. Data are representative of at least three independent experiments. **B and C**. Growth of PA14 and PAO1 strains listed in M63 liquid with ethanol as the sole carbon source in 21% oxygen. Statistics based on one-way ANOVA and Tukey’s multiple comparison; between group comparisons were significant with a *P*<0.001 in panel B and *P*<0.0001 in panel C.

**Table S1.**
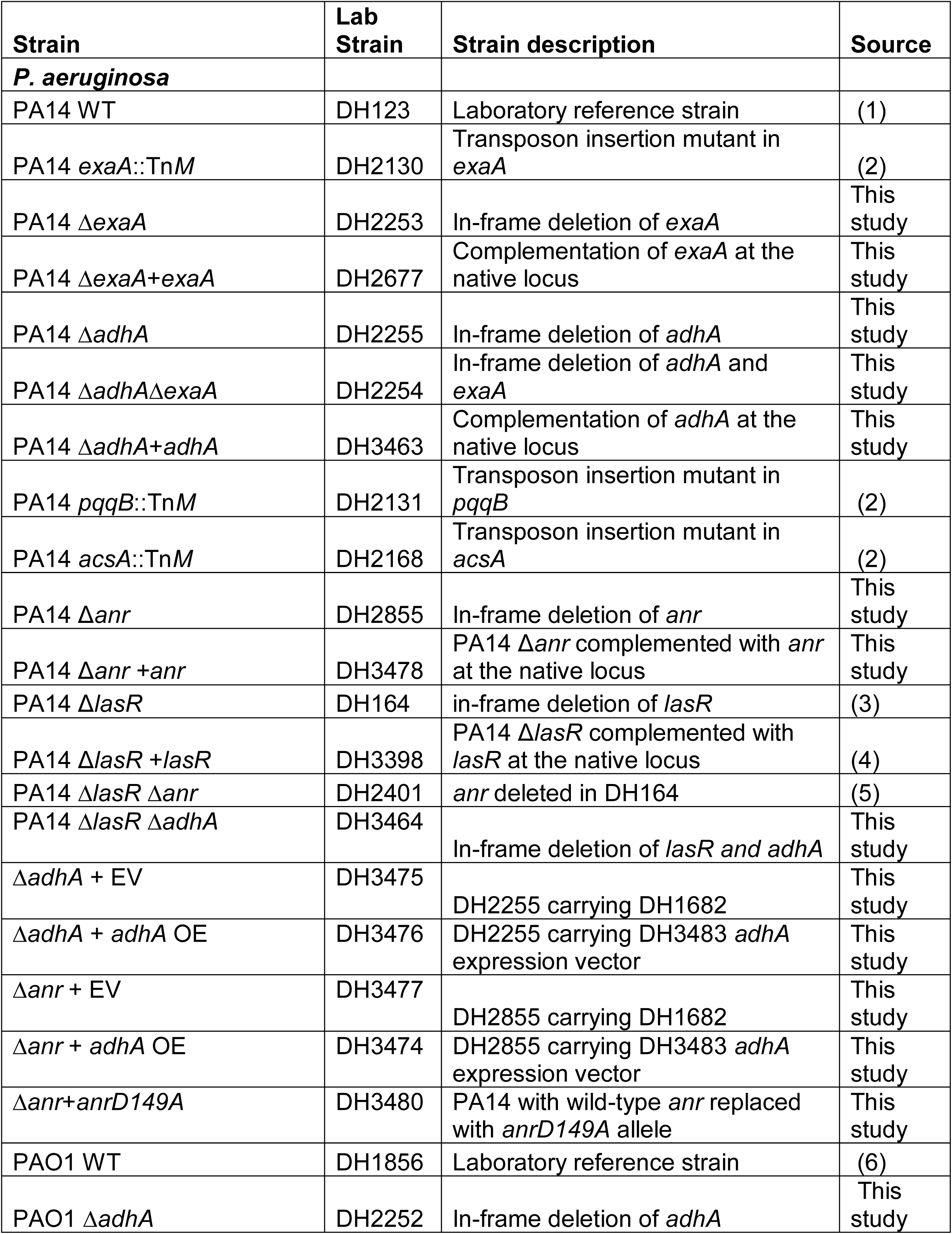
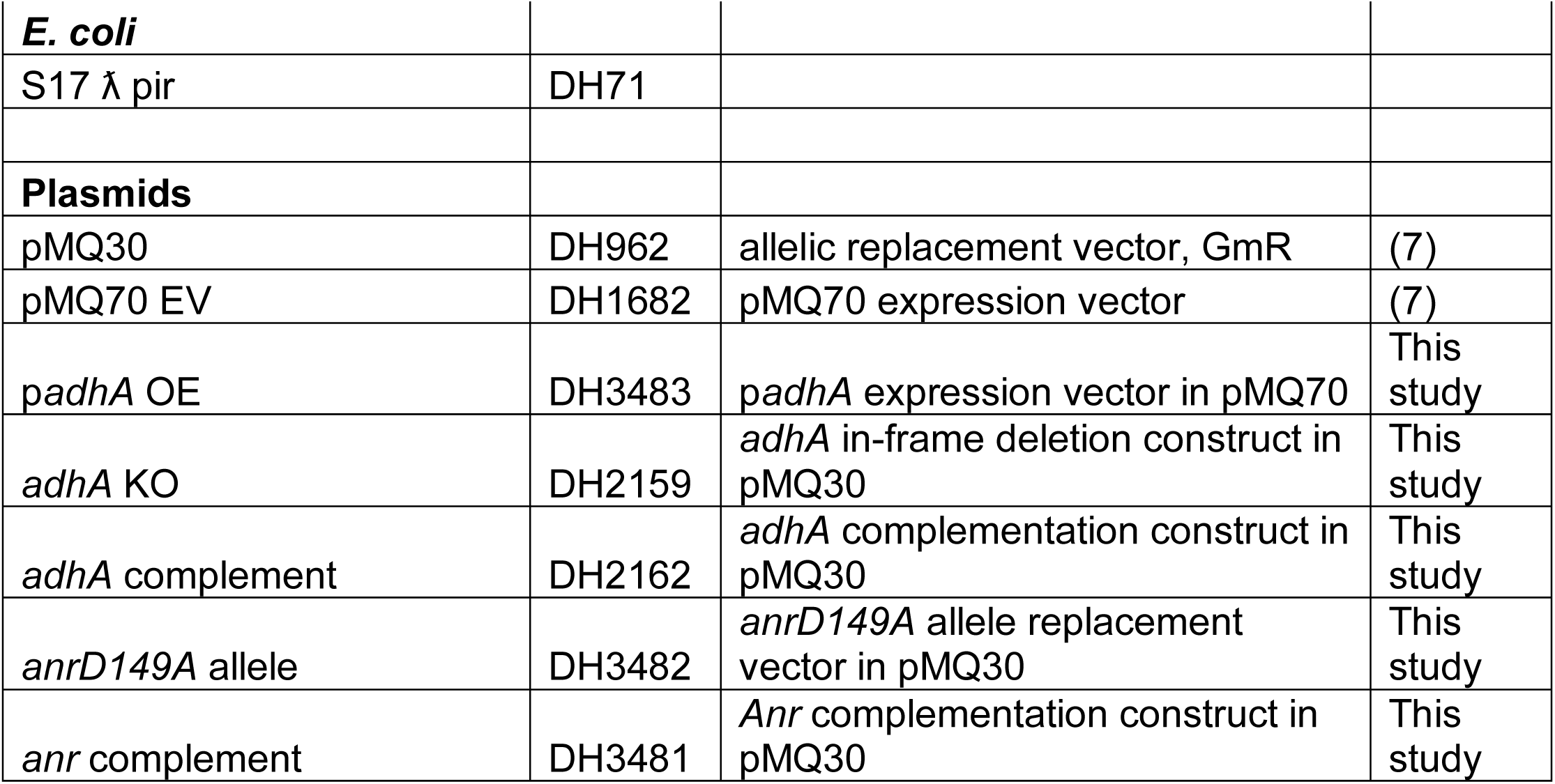
Strains used in this study.

**Table S2.**
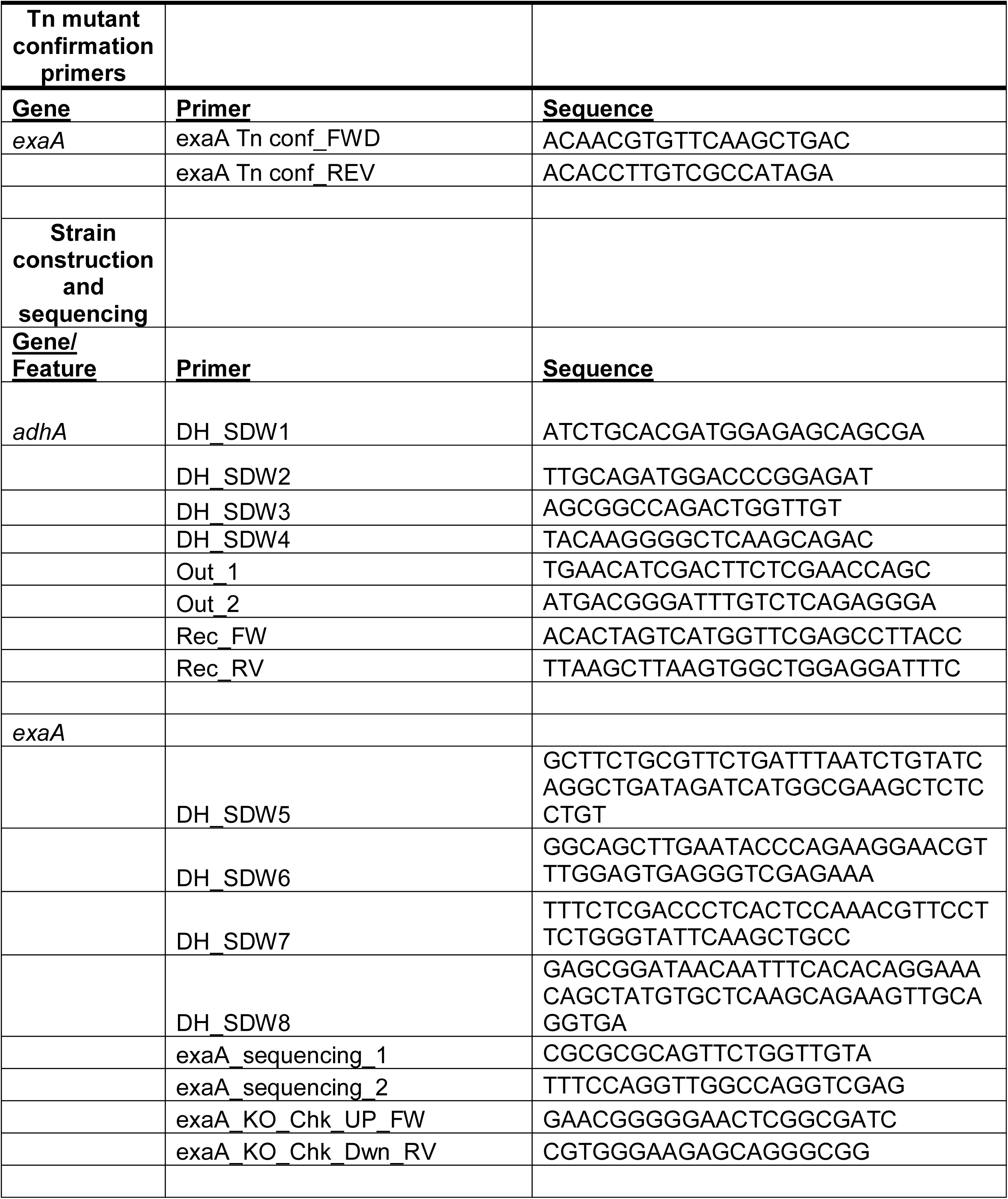
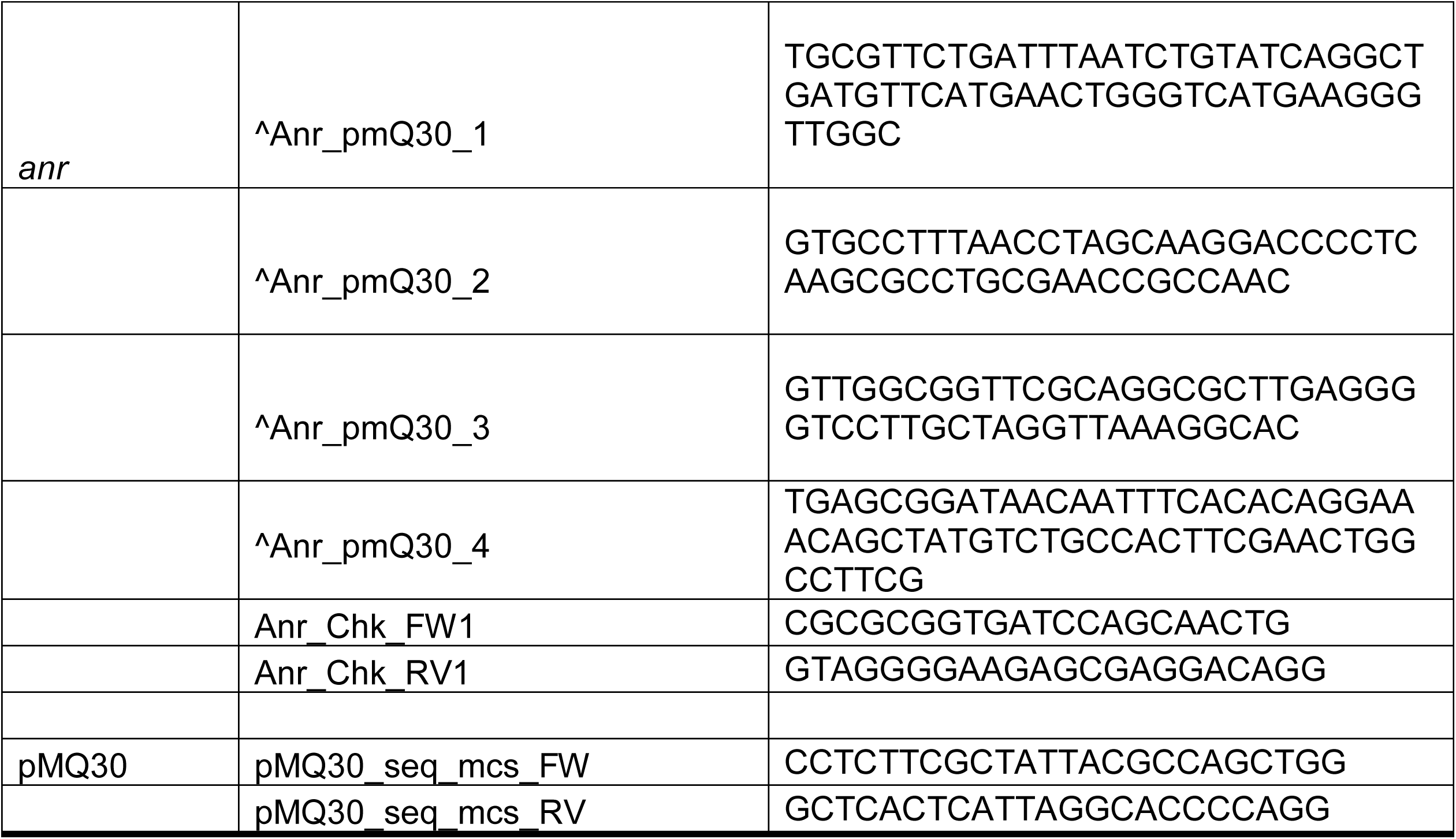
Primers used in this study.

